# Insights from spatial measures of intolerance to identify pathogenic variants in developmental and epileptic encephalopathies

**DOI:** 10.1101/2021.07.28.454182

**Authors:** M Silk, A De Sá, M Olshansky, D B Ascher

## Abstract

Developmental and epileptic encephalopathies (DEEs) are a group of epilepsies with early onset and severe symptoms that sometimes lead to death. While a number of genes have been successfully implicated, it remains challenging to identify causative mutations within these genes from the background variation present in all individuals due to disease heterogeneity. Our ability to detect likely pathogenic variants has continued to improve as *in silico* predictors of deleteriousness have advanced. We investigate their use in prioritising likely pathogenic variants in epileptic encephalopathy patient whole exome sequences and show that the inclusion of structure-based predictors of intolerance improve upon previous attempts to demonstrate enrichment within epilepsy genes.

## INTRODUCTION

Developmental and epileptic encephalopathies (DEEs) are a class of severe, early-onset epilepsies characterised by hypsarrhythmia detected by EEG, infantile spasms, multi-form seizures, cognitive and behavioural deficits, and can sometimes lead to death. Many studies have investigated a likely genetic basis underpinning the disease, however identifying causal variants in patients remains challenging due to the apparent disease heterogeneity^1^.

Studies investigating the genetic basis of DEE have primarily focused on *de novo* mutations (DNMs) as, due to the early onset and severity of the disease, few causal variants are disseminated and maintained in the healthy population. Causative variants have been identified in 30-50% of infants with severe DEE, however in many cases multiple DNMs may be identified with no prior history of functional studies or previous observation in epilepsy patients^2^. To prioritise candidate DNMs likely causative of a patient's phenotype, *in silico* predictors of deleteriousness have proven to be beneficial, however their accuracy remains imperfect.

A previous study (Petrovski et al., 2014)^3^ aimed to identify genes linked to DEE and to identify causal DNMs in a cohort of 356 patients through a collaboration between two consortia (EuroEPINOMICS and Epi4K/EPGP)^4^. A likelihood analysis revealed an enrichment of DNMs in patients compared with a cohort of 411 trio exome sequencing controls. 75% of the 429 DNMs identified in patients were predicted to disrupt synaptic transmission regulation.

Since this analysis, our repertoire of useful datasets and tools have grown, which can greatly improve our ability to assess a variant's likelihood of pathogenicity. Recent large datasets of population variation such as gnomAD^5^ allow us to filter observed variants in patients that are present in the healthy population. Recent measurements of missense intolerance in particular can be highly informative for DNM assessment, where it is likely that pathogenic DNM’s cluster within intolerant regions due to their severity^6^.

We revisit the missense variants in this cohort to identify whether there is significant enrichment of predicted damaging variants in patient samples when compared with controls, and using recent *in silico* predictors of pathogenicity, we investigate whether these new tools allow us to identify a greater number of causative variants or epilepsy-associated genes.

Additionally, we investigate the enrichment of likely pathogenic missense variants within genes associated with DEE within the publically available Epi25K dataset. Our analysis shows an enrichment of these variants within these genes despite including both *de novo* and non *de novo* variants within the DEE patient whole exome sequencing experiments.

## RESULTS

### Pathogenic and background missense variants among epilepsy genes

We examined missense variants exome-wide and within 34 genes implicated in epileptic encephalopathies using the Epi4K case-ascertained *de novo* missense variants, Epi25K case-ascertained missense variants and ClinVar missense variants (accessed 16th June 2020).

34 key genes implicated in epileptic encephalopathies were selected based on a comprehensive literature review by He et al. (2019)^7^. Of the 34 genes, 14 are components of ion channels (SCN1A, SCN2A, SCN8A, KCNA1, KCNA2, KCNB1, KCNQ2, KCNT1, CACNA1A, GRIN1, GRIN2A, GRIN2B, GRIN2D, HCN1) and 20 are not (ATP1A2, NTRK2, SLC2A1, SLC6A1, STXBP1, FGF12, YWHAG, DYNC1H1, SPTAN1, ANKRD11, EEF1A2, FOXG1, NACC1, CHD2, DNM1, DNM1L, GNAO1, HECW2, NEDD4L, SYNGAP1).

We evaluated the distributions of commonly utilitised sequence-based *in silico* predictors of deleteriousness on case-ascertained and control variants both exome-wide and within these 34 genes. 276 case-ascertained confirmed *de novo* variants from the Epi4K consortium dataset were compared with 454 control *de novo* variants not derived from epilepsy patients (control group 1). A second set of 762 control variants were used for comparison from studies of autism spectrum disorders (control group 2). All variants were annotated using the Variant Effect Predictor (release 101) and the plugin dbNSFP 3.5a, confirming that all are missense variants. 1,082,844 missense variants from the Epi25K consortium dataset’s developmental and epileptice encephalopathies (DEE) analysis group were also included for comparison. These were compared with the DiscovEHR missense variants, derived from the general population and presumed to be largely depleted of pathogenic variation. We considered only variants in DiscovEHR not present in the Epi25K dataset.

Of the 276 case-ascertained *de novo* missense variants, 29 variants were located within 15 of the 34 genes implicated in DEE (CACNA1A: 1, DNM1: 5, GNA01: 2, GRIN1: 1, GRIN2B: 1, HECW2: 1, KCNB1: 1, KCNQ2: 2, KCNT1: 1, NEDD4L: 1, SCN1A: 4, SCN2A: 2, SCN8A: 2, STXBP1: 4, YWHAG: 1), compared with only 1 control group 1 variant (NTRK2: 1) and 3 from control group 2 (GRIN1: 1, SCN1A: 1, SLC2A: 1). These are shown in Figure 1 as lollipop plots^8^.

To assess the predictive power of a range of *in silico* predictors, the ClinVar variant summary dataset was used. For accurate comparison to the other variant sets under investigation, these were annotated using the same approach and filtered to missense variants. Variants with unknown or conflicting significance were removed. While not all variants in this set have been confirmed to be related to DEE, the predictive tools used in this study can still provide insight to likely impacts on protein function regardless of the resulting phenotype.

**Figure 1:**
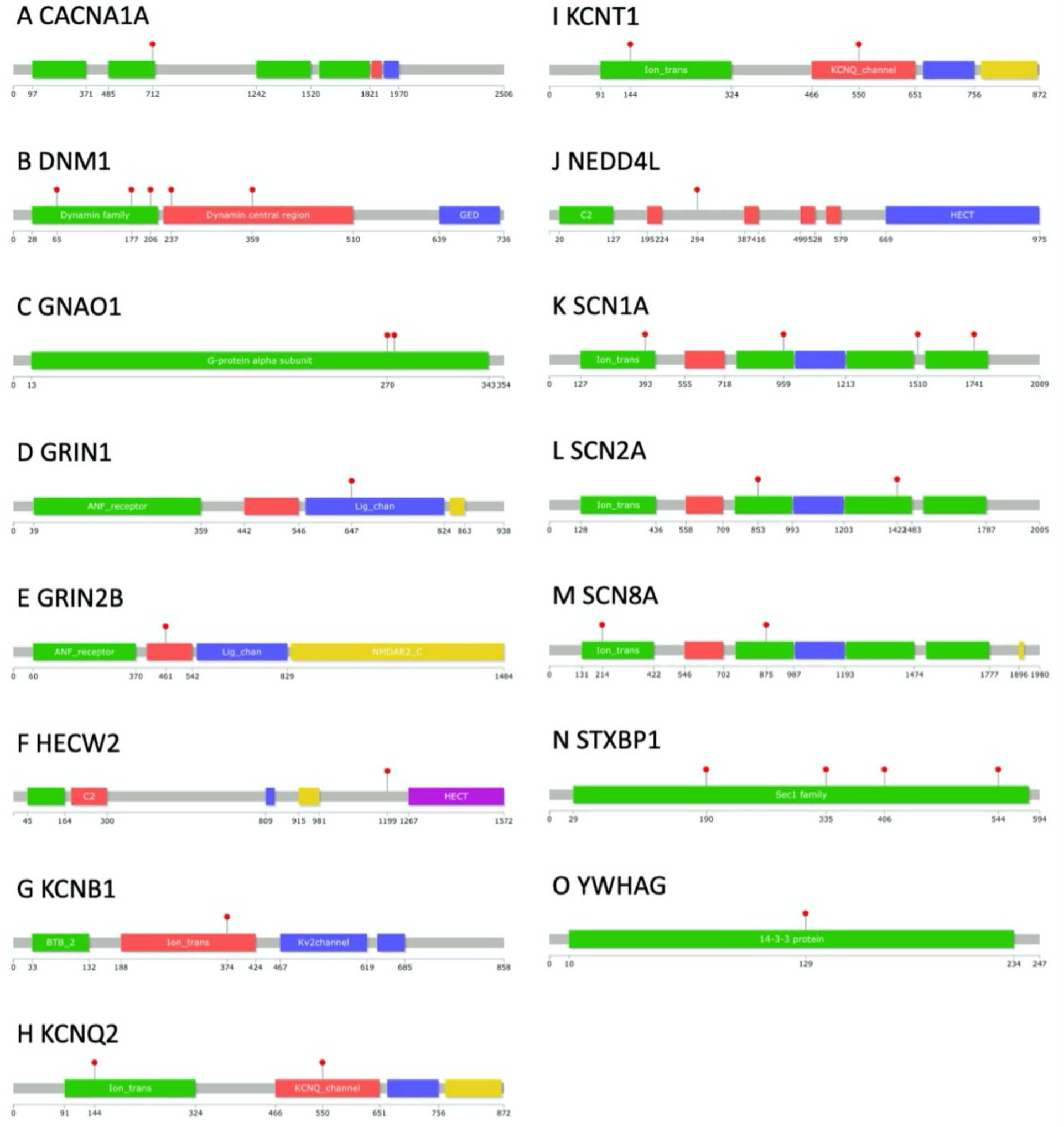
Case-ascertained *de novo* mutation locations within epilepsy genes. Red lollipops denote locations of epilepsy variants, identified in 15 of the 34 target genes. Domains are sourced from Pfam.

### Identifying missense-intolerant regions in epilepsy genes

We previously investigated missense intolerance within epilepsy genes, and have shown that the Missense Tolerance Ratio (MTR) can be a powerful tool to identify functionally important regions within genes^9^. Since this study, the MTR has been recalculated using additional variation from gnomAD v2 including exomes and genomes, UK Biobank’s 50,000 exomes and DiscovEHR, increasing the accuracy of scores.

We examined the predictive utility of the updated MTR scores by comparing the number of unique Epi4K variants within the top 25% most intolerant MTR scores exome-wide, corresponding to an MTR score < 0.78 with control group sets. Of the 276 case-ascertained variants, we observed 91 (33%) within highly intolerant regions compared with 88 (19%) control group 1 variants, and 147 (19%) control group 2 variants, indicating that other genes harboring *de novo* variants may also be implicated in disease outcomes.

Considering only variants within the 34 EE genes, 21 (72%) of case-ascertained variants were found within intolerant regions, as well as the single (100%) control group 1 variant and 2 (50%) control group 2 variants. While we lack sufficient sample sizes to draw direct comparisons between these, it is unsurprising that many control variants are within intolerant regions given how intolerant and conserved these genes are.

Additionally, we examined whether there is enrichment of Epi25K missense variants within the 34 DEE genes compared with the DiscovEHR control variants, after filtering any Epi25K missense variants from the DiscovEHR control variants. 4656 of the total 10,740 missense variants (43%) were observed within intolerant regions, compared with 6,867 of 17,229 DiscovEHR missense variants (40%).

### Investigating spatial missense intolerance within protein structures of epilepsy genes

We next investigated intolerance within the 34 EE genes in the context of their protein tertiary structures, utilising experimentally determined structures from the RCSB PDB^10^ and homology modelled structures from SWISS-MODEL^11^. 115 case-ascertained variants, 170 control group 1 variants and 263 control group 2 variants had a valid MTR3D score for analysis. Defining MTR3D < 0.75 as intolerant, we observe 47 case-ascertained variants (41%), 36 control group 1 variants (21%) and 58 control group 2 variants (22%). Delineating MTR3D < 0.5 as strongly intolerant, we observe 22 (19%) case-ascertained variants, 7 control group 1 (4%) and 14 control group 2 missense variants (5%). Thus, we observe significant enrichment of *de novo* mutations within intolerant regions in the epilepsy analysis group compared with both control groups across all genes with observed DNMs, suggesting that other genes are likely implicated in disease outcomes.

Specific to the 34 EE genes, we were able to derive MTR3D scores for 17 case-ascertained variants only. 16 of the 17 had an MTR3D score below 0.75, and 10 of which had an MTR3D score below 0.5 denoting strong intolerance.

Additionally, we utilised the MTRX, a combined measure of intolerance built through a Random Forest approach using the MTR v1 (41 codons), MTR v2 (21 codons), MTR3D and residue solvent accessibility (RSA). MTRX scores were available for 92 case-ascertained variants, 151 control group 1 variants and 238 control group 2 variants. MTRX scores approaching 1 are considered more likely deleterious. Using a cutoff of 0.75, we observe 38 case-ascertained (41%), 28 control group 1 (19%) and 60 control group 2 variants (25%) within these intolerant regions. 12 case-ascertained variants were located within the 34 EE genes with a valid MTRX score, 11 of which were within intolerant regions, with an overall mean MTRX score of 0.93.

Next, we calculated conservation for all genes and compared this with the MTR estimates. Considering all *de novo* variants, a Pearson’s correlation of 0.03 (p < 0.58) was observed, indicating low correlation between these scores. However, in the 34 DEE genes, we observe a Pearson’s correlation of 0.34 (p < 0.001) indicating that they are moderately correlated within these genes.

### Evaluating the predictive utility of in silico predictors of deleteriousness

*In silico* predictors of deleteriousness are of high value in prioritising likely deleterious variants from among background variants when diagnosing rare disease. To assess their efficacy in epilepsy, we compared the predictions of a number of scores obtained through dbNSFP between the case-ascertained DNM variants from Epi4K and the control groups as described above.

Variants were first examined to identify whether the distribution of case-ascertained DNMs feature significantly different scores compared to control variants. Currently, there are over 100 different *in silico* predictors available, however for the purposes of this study we selected a subset that are available in Variant Effect Predictor (VEP) and through dbNSFP. This set provides a mixture of scores created using physicochemical properties, standing variation in the human population and combined approaches through machine learning. We utilised the rank scores provided for each metric, where each score is converted to a percentile based on all scored positions within the dataset.

Owing to the small number of control *de novo* variants observed within the 34 EE genes, we compared *in silico* predictions for these as a combined set.

We first examined Pearson’s correlation between each of the predictive tools’ rank scores (Figure 2). Correlations were found to be overall higher for the subset of variants within the DEE genes. Overall scores were found to be somewhat correlated across all genes, especially those derived from similar properties.

**Figure 2:**
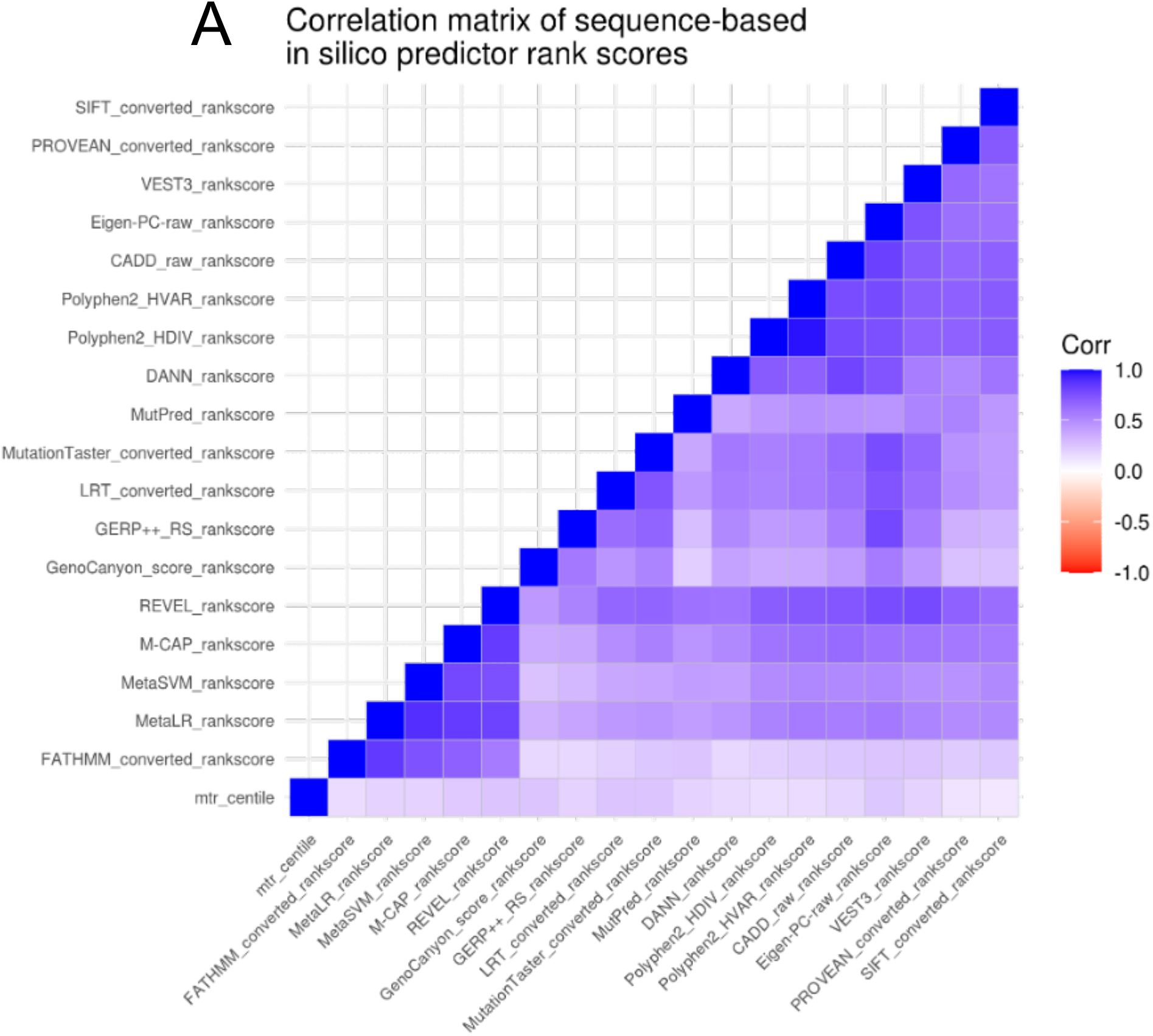
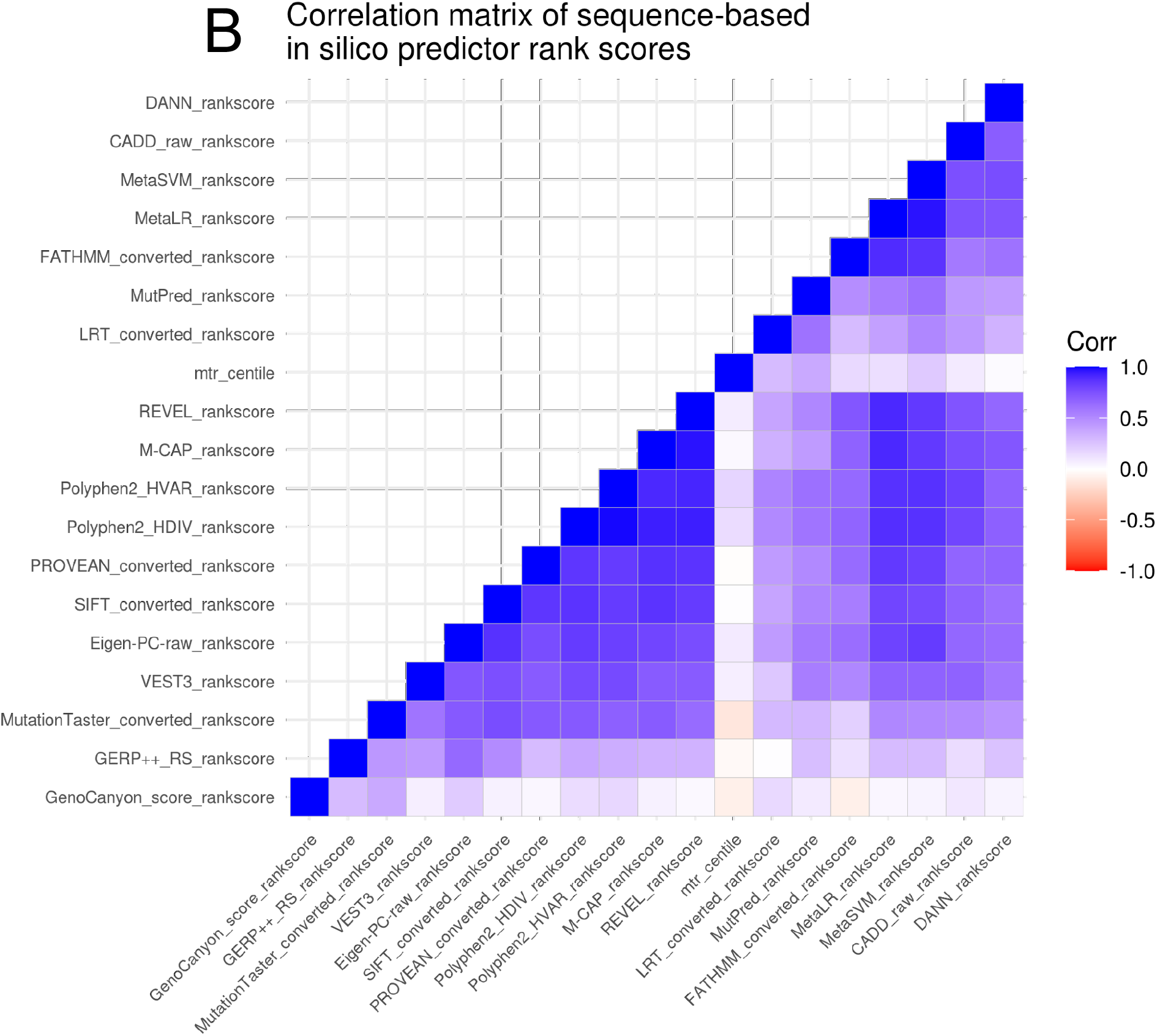
Correlation plot of *in silico* predictor scores for complete cases of variants. (A) Correlation of scores for all Epi4K variants across all genes. (B) Correlation of scores for Epi4K variants within the 34 EE genes.

Wilcox signed rank tests comparing epilepsy DNMs to control groups identified a significant difference in rank scores for CADD (p < 0.008), DANN (0.03), Eigen-PC-raw (0.0008), FATHMM (p < 0.002), GenoCanyon (p < 0.02), LRT (p < 0.01), M-CAP (p < 3.8 * 10-5), MetaLR (p < 0.0004), MetaSVM (p < 0.0003), MutPred (p < 0.02), MutationTaster (p < 0.01), PROVEAN (p < 5.3 * 10-5), Polyphen2-HDIV (p < 2.9 * 10-5), REVEL (p < 1.2 * 10-6), SIFT (p < 0.002) and VEST3 (p < 3.8 * 10-6) and MTR (p < 1.3 * 10-6). We also note scores in GERP++ RS differ but did not achieve significance (p < 0.06). The distributions of these scores are shown in Figure 3.

**Figure 3:**
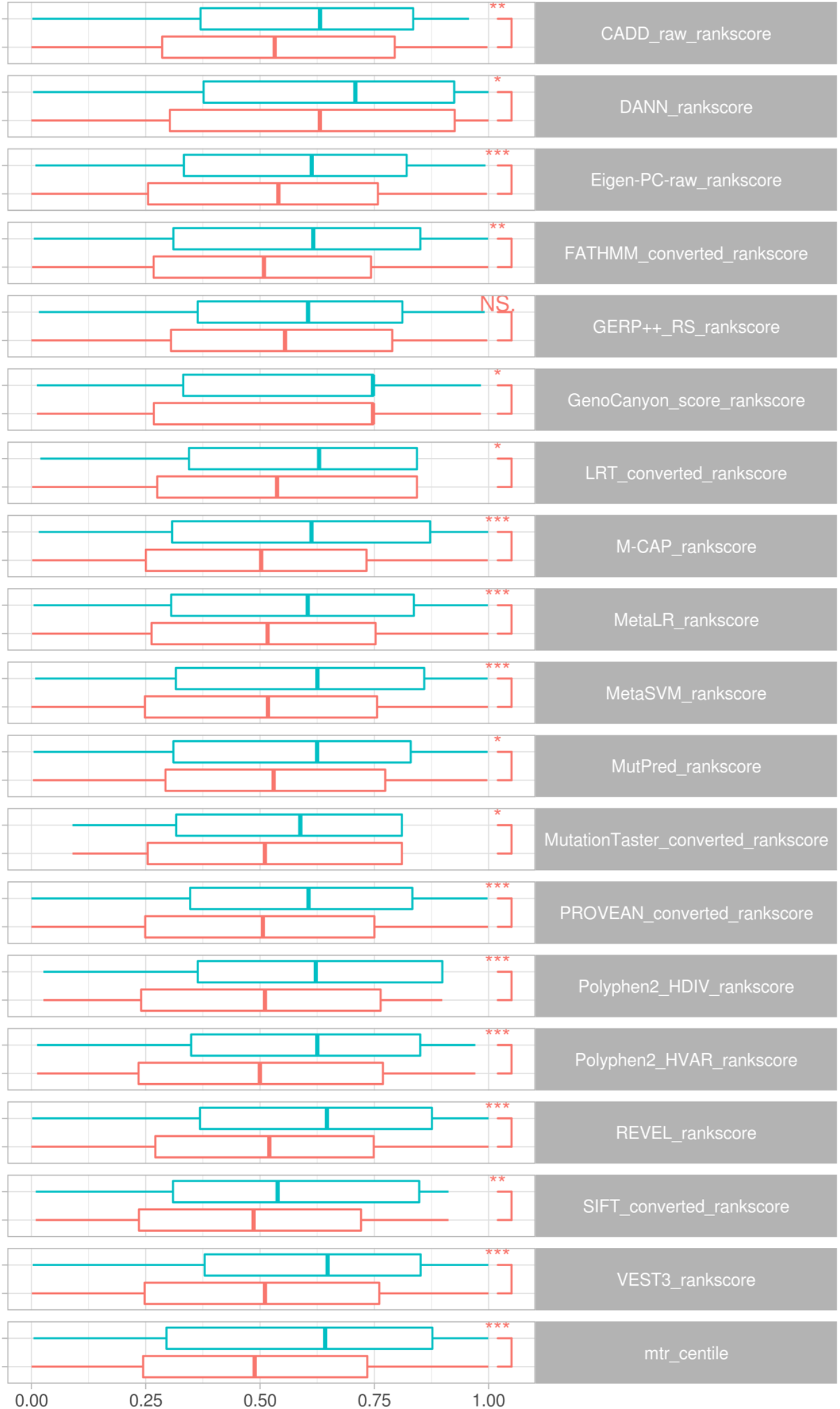
Comparison of *in silico* predictors of deleteriousness. Boxplots display rank score distributions for *de novo* missense variants across all genes, comparing case-ascertained variants (salmon) with combined variants from control group 1 and 2 (blue).

Examining the variants specific to the DEE genes, we see similar separation between case-ascertained and control variants for CADD (p < 0.005), DANN (p < 0.02), Eigen-PC-raw (p < 0.003), LRT (p < 0.007), M-CAP (p < 0.05), MetaLR (p < 0.01), MutationTaster (p < 0.0001), PROVEAN (p < 0.01), Polyphen2-HDIV (p < 5.5 * 10-5), REVEL (p < 0.009), SIFT (p < 0.01) and VEST3 (p < 0.004). Statistical significance was not achieved for FATHMM (p < 0.08) and GERP++ RS (p < 0.12), GenoCanyon (p < 0.11), MetaSVM (p < 0.15), MutPred (p < 0.06).

## DISCUSSION

*In silico* predictors of deleteriousness continue to show great utility in identifying likely pathogenic variants, with key relevance to developmental and epileptic encephalopathies where the disease has been shown to often be attributed to a single variant. Our repertoire of tools continues to expand as newer approaches are conceptualised and novel biological datasets are made available through which meaning can be discovered for each variant.

Difficulties arise where these tools are built using different human reference genome builds and different transcript versions. The Variant Effect Predictor and ANNOVAR have proven to be extremely useful in this regard, however issues are inevitable as each predictor is built and updated from differing transcripts.

Intolerance-based predictors and conservation were shown to be highly effective predictors, owing to the severity and early onset of the disease. As shown in a previous study of ClinVar variants, we have observed this to be the case in genes with known dominant and recessive patterns of inheritance^12^, with a greater enrichment of pathogenic variants in dominant genes. Of the 34 DEE genes under investigation, 30 are within the top 10% most intolerant genes indicating that variants arising within these genes in most regions (and not just the known domains) are likely deleterious. Despite the overall strong intolerance of these genes, we still see significant differences in the MTR predictions between the case and control *de novo* variants.

Utilising the spatial-based MTR3D where a successful alignment between sequence and structure was available and a score could be derived, we identified 16 of 17 case-ascertained *de novo* variants residing within intolerant - scored regions. Due to few control variants residing within the 34 DEE genes and with none located in a region with a valid MTR3D score, it is challenging to draw conclusions from this result. Similarly, this remains a challenge for many structure-based predictors that rely on the availability of a resolved or homology modelled protein structure.

Where these can be calculated however, we see great potential in their utility as an additional tool for variant prioritisation. It is clear that the accuracy of predictive tools as a whole continue to improve. It is particularly surprising that *de novo* missense variants exterior to DEE genes were deemed to be more likely deleterious than control *de novo* missense variants.

These results confirm the utility of *in silico* predictors for prioritisation of variants in DEE and suggest that additional genes are of interest to further our understanding of the disease.

## METHODOLOGY

### Study subjects and sequencing procedures

Three epileptic encephalopathy cohorts were evaluated for this study from the Epi4K *de novo* mutations study: (1) Epilepsy Phenome/Genome Project cohort 1 (n = 264 trios), (2) Epilepsy Phenome/Genome Project cohort 2 (n = 73 trios), and (3) EuroEPINOMICS-RES cohort (n = 19 trios). Informed consent was obtained from parents or legal guardian of each participant, and studies were approved by local ethics committees of each participating centre. Further information can be obtained from the Epi4K publication^4^.

Missense variants from the Epi25K whole-exome sequencing variant-level summary were also included for analysis, filtered to only developmental and epileptic encephalopathy patients (n = 1,021), thus removing samples from genetic generalised epilepsy (n = 3,108) and non-acquired focal epilepsy (n = 3,597). This dataset was downloaded from the Epi25 WES Browser and is publically available pre-filtered for missense variants with MPC >= 2, including both *de novo* and non-*de novo* variants.

The DiscovEHR whole-exome sequencing dataset was used to compare with the epilepsy cohorts^13^, including exomes from 50,000 individuals.

### Missense variant annotation

Epi4K *de novo* variants were annotated using the Variant Effect Predictor (version 101)^14^ and the dbNSFP 3.5a plugin^15^ to include a range of predictors of deleteriousness based on physicochemical properties, conservation and combined approaches. dbNSFP’s rank scores were also included, where each score is given a ranking from 0-1 based on its percentile across all scored coding positions within the dbNSFP dataset,, where scores closer to 1 are considered more likely severe. Following annotation, unique variants classified as missense were selected for subsequent analysis (Case-ascertained variants: N = 276. Control group: N = 454).

The same annotation procedure was applied to the Epi25K missense variants (N = 1,082,844) and the DiscovEHR control population variants (N = 2,134,301). Similarly, these sets are filtered to only those with missense consequences.

Variants were further annotated with Missense Tolerance Ratio (MTR), Missense Tolerance Ratio-3D (MTR3D) and Missense Tolerance Ratio consensus (MTRX) scores to explore regional intolerance to missense variation at each variant’s position. MTR3D and MTRX scores are available for a subset of epilepsy-related genes depending on availability of protein tertiary structures.

A subset of the Epi4K *de novo* missense variants was selected where these reside within 34 genes with evidence of being implicated in developmental and epileptic encephalopathies (DEE) as described by He et al. (2019)^7^.

ClinVar variants were annotated using the same procedure described above and filtered to those predicted to be missense and benign, likely benign, pathogenic or likely pathogenic, omitting those with unknown significance^16^. To examine the distribution of pathogenic variants within epilepsy genes, these were subset to the previously described 34 epileptic encephalopathy genes. To ensure that these are mutually exclusive to test sets, variants were filtered to exclude 16 variants overlapping with the case-ascertained *de novo* variants.

### Calculating conservation

We utilised the toolset by Capra and Singh (2007)^17^ to estimate sequence conservation based on Jensen-Shannon divergence from a multiple sequence alignment. Some positions where variants were located could not be measured due to gaps in aligned sequences lowering confidence in the results.

### Annotating variants with MTR3D scores

As the MTR3D predicted scores include mappings between sequence protein positions and structural residue numbers, these were used to map the Epi4K *de novo* variants, Epi25K variants and ClinVar pathogenic and benign variants to protein structures. An experimentally resolved structure from the RCSB Protein Data Bank was used preferentially where MTR3D mappings were available, and otherwise a homology modelled protein tertiary structure was used from the SWISS-MODEL database. Where multiple structures exist where a variant could be mapped, the structure with the highest proportion of matching sequence and structural positions was selected. This varies due to partial experimental structures representing only certain regions and domains of a gene, residues that may be different due to different transcripts or substitutions to assist in the stability of the structure’s creation. Multiple structures were used for genes where partial structures best fit for different variants (see Supplementary Table S1).

