## Supplemental Data for "Insights from spatial measures of intolerance to identify pathogenic variants in developmental and epileptic encephalopathies"

### Supplementary Table S1: Structures used for MTR3D calculations

| HGNC Symbol | PDB ID | Source |
|---|---|---|
| ANKRD11 | 99_339_6by9.1.A_5f76d4ace473290e766cc658.pdb | SWISS-MODEL |
| ATP1A2 | 26_1020_4ret.2.A_5f7050908ae8c48a4ab2c2fc.pdb | SWISS-MODEL |
| CACNA1A | 1232_1947_5gjv.1.A_5cc87686732c1ef8ec7488d8.pdb | SWISS-MODEL |
| CACNA1A | 3BXK | Experimentally determined |
| CHD2 | 259_1022_3mwy.1.A_5f68031dc60d5f991ca20e5a.pdb | SWISS-MODEL |
| CHD2 | 306_990_6ftx.1.M_5f68031dc60d5f991ca20e4a.pdb | SWISS-MODEL |
| DNM1 | 6DLU | Experimentally determined |
| DNM1L | 4BEJ | Experimentally determined |
| DNM1L | 1_736_5wp9.1.A_5f67078e80434655032fe539.pdb | SWISS-MODEL |
| DYNC1H1 | 5NUG | Experimentally determined |
| DYNC1H1 | 1444_4646_5nug.1.A_5f686bb9d903f866a892fff7.pdb | SWISS-MODEL |
| EEF1A2 | 4_461_6ra9.1.B_5f653e249950e55facbb0951.pdb | SWISS-MODEL |

| | | |
|---|---|---|
| FGF12 | 4JQ0 | Experimentally determined |
| FOXG1 | 184_289_6qvw.1.A_5f65713f55ba95672fbdf47a.pdb | SWISS-MODEL |
| GNAO1 | 6OIK | Experimentally determined |
| GNAO1 | 12_349_4gnk.1.A_5f6440dfb61a4ebe2dc96a6f.pdb | SWISS-MODEL |
| GRIN1 | 6IRA | Experimentally determined |
| GRIN1 | 25_841_6whs.1.A_5f7061b38ae8c48a4ab396f8.pdb | SWISS-MODEL |
| GRIN2A | 6IRH | Experimentally determined |
| GRIN2A | 32_840_5iou.1.D_5f705ad68ae8c48a4ab33f3e.pdb | SWISS-MODEL |
| GRIN2B | 5EWM | Experimentally determined |
| GRIN2B | 34_845_6whs.1.B_5f67d87fadf0a0f8fbc5565a.pdb | SWISS-MODEL |
| GRIN2D | 49_870_5ide.1.C_5f704e0e8ae8c48a4ab29e27.pdb | SWISS-MODEL |
| HCN1 | 6UQF | Experimentally determined |
| HCN1 | 94_633_6uqf.1.A_5f7050f98ae8c48a4ab2cb0b.pdb | SWISS-MODEL |
| HECW2 | 2LFE | Experimentally determined |
| HECW2 | 1181_1566_5tj7.4.A_5f67e458d6c129eb4b1cdefc.pdb | SWISS-MODEL |
| KCNA1 | 32_419_5wie.1.B_5f70502e8ae8c48a4ab2c071.pdb | SWISS-MODEL |
| KCNA2 | 31_421_6ebk.1.B_5f6588141a1d1f370635a17f.pdb | SWISS-MODEL |
| KCNB1 | 29_424_6ebk.1.B_5f66f816b078d19d780f175d.pdb | SWISS-MODEL |
| KCNQ2 | 6FEG | Experimentally determined |
| KCNQ2 | 70_329_7cr1.1.A_5f6d90cc545f5a798b410058.pdb | SWISS-MODEL |
| KCNT1 | 71_1200_5u70.1.D_5f7f1ef24ec6d929eacf0c96.pdb | SWISS-MODEL |
| NACC1 | 4U2N | Experimentally determined |
| NACC1 | 3_124_6w66.1.C_5f65b96d4df9e3a0090caa39.pdb | SWISS-MODEL |

| | | |
|---|---|---|
| NEDD4L | 384_969_5xmc.1.A_5f673b25e33774264de5fbed.pdb | SWISS-MODEL |
| NEDD4L | 3JVZ | Experimentally determined |
| NTRK2 | 4AT5 | Experimentally determined |
| SCN1A | 1790_1942_4dck.1.A_5f6816acab0d00685490c2f3.pdb | SWISS-MODEL |
| SCN2A | 6J8E | Experimentally determined |
| SCN2A | 1788_1929_4jpz.1.B_5e1273f59ffd12a1ae2ee8f2.pdb | SWISS-MODEL |
| SCN8A | 1778_1921_4jpz.1.B_5f68131a033d6797c5fb9d45.pdb | SWISS-MODEL |
| SLC2A1 | 4PYP | Experimentally determined |
| SLC6A1 | 42_576_4xp4.1.A_5f661e539172bfe4f51b18c8.pdb | SWISS-MODEL |
| SPTAN1 | 3FB2 | Experimentally determined |
| SPTAN1 | 1828_2472_4d1e.1.B_5f6838d52a32a9d4f0792f88.pdb | SWISS-MODEL |
| STXBP1 | 3_592_4jeu.1.A_5f6617f21ac1d9a920966478.pdb | SWISS-MODEL |
| SYNGAP1 | 252_733_3bxj.2.A_5f70515f8ae8c48a4ab2cf8e.pdb | SWISS-MODEL |
| YWHAG | 6BYJ | Experimentally determined |
